# Differential requirement of *nanos* homologs in the germline suggests the evolutionary path toward an inheritance mechanism of primordial germ cell formation in the silkmoth *Bombyx*

**DOI:** 10.1101/2022.11.21.517435

**Authors:** Hajime Nakao, Yoko Takasu

**Affiliations:** Insect Design Technology group, Division of Insect Advanced Technology, Institute of Agrobiological Sciences, National Agriculture and Food Research Organization (NARO), 1-2 Oowashi, Tsukuba, Ibaraki 305-8634, Japan; Silk Materials Research Group, Division of Silk-Producing Insect Biotechnology, Institute of Agrobiological Sciences, National Agriculture and Food Research Organization (NARO), 1-2 Oowashi, Tsukuba, Ibaraki 305-8634, Japan

**Keywords:** *Bombyx*, lepidopteran insect, PGC specification, *nanos*, segmentation

## Abstract

The lepidopteran insect *Bombyx mori* possesses unique embryogenesis characteristics among insects. *nanos* (*nos*) has conserved functions in metazoan primordial germ cell formation. *Bombyx* possesses four *nos* genes (M, N, O, P), a unique feature found in lepidopterans examined so far. Of these, maternal *nosO* mRNA exhibits a localization pattern: it may act as a primordial germ cell (PGC) determinant. A previous knock-out experiment of *nosO* showed that this localized mRNA is dispensable for PGC formation in laboratory environment and has limited involvement in PGC specification. This study examined whether other nos genes act redundantly with *nosO* in germline using RNAi and gene editing. Although individual embryonic RNAi exhibited no detectable phenotypic alterations, simultaneous RNAi of *nosO*/ *nosP* markedly reduced oocyte number and male fecundity. Additionally, *nosP* KO almost completely sterilized both sexes. Because *nosP* is broadly expressed in the posterior of embryos in non-germline specific manner, these results could reflect an evolutionary step taken by *Bombyx* toward its unique inheritance mechanisms. This study also suggests that *nos* genes in *Bombyx* do not affect anterior-posterior axis specification. This could reflect its characteristic embryogenesis.

## Introduction

During insect embryogenesis, diverse mechanisms using similar sets of genes are employed to converge on a similar body plan exemplified by the state achieved at the phylotypic stage. How environment and developmental systems interact to produce such diversity is of interest (Church *et al*., 2019; Dixon, 1994; Evans, 2014; Whittle and Extavour, 2016, 2017). The lepidopteran insect, *B. mori* is an interesting organism to study because it is phylogenetically close to *Drosophila*. Yet, the early embryonic processes are quite distinct among insects (Nakao, 2021).

In the past decades, efforts have been made to describe such diversity in molecular terms across insect orders, notably in segmentation and germ cell specification mechanisms (Davis and Patel, 2002; Extavour and Akam, 2003; Peel *et al*., 2005; Lynch, 2014; Quan and Lynch, 2016).

*Drosophila* germ cell specification and PGC formation mechanisms are the best examples, where the localized maternal cytoplasmic determinant called pole plasm at the posterior egg pole dictates PGC cell fate to the cells that sequester the determinant during cellularization. This is an example of inheritance mechanisms of PGC formation, which is widely distributed in the animal kingdom, including vertebrates. The inherited cytoplasm is generally known as germplasm (Eddy, 1976). Zygotic induction is another PGC formation mechanism well-known in studies in mice, but is also known in insects. Zygotic induction is considered an evolutionary ancient mechanism from which inheritance mechanisms are derived. The distinction is classically well-known, and relatively recently, the molecular underpinnings have begun to be uncovered (also see below). *vasa* (*vas*) and *nanos* (*nos*) are classic examples of germ cell marker genes, the products of which are often associated with the germplasm (Extavour and Akam, 2003; Quan and Lynch, 2016). *nos*, which encodes a zinc finger translational repressor protein, is found to be essential for the formation/maintenance of germ cells. In PGCs, it suppresses somatic developmental programs, thus preserving the undifferentiated state required for germ cell function (Kobayashi *et al*., 1996; De Keuckelaere *et al*., 2018).

In insect, *nos* involvement in embryo segmentation is also well-known, and this function may be evolutionarily conserved (Lall *et al*., 2003). In *Drosophila*, segmentation occurs nearly simultaneously through a series of hierarchical genetic cascades that begins with molecular cues emanating from the anterior and posterior poles of egg cytoplasm after fertilization leading to the establishment of the positional information along the anterior–posterior axis. *nos* contributes to this positional information establishment process by suppressing initially uniformly distributed *hunchback* (*hb*) mRNA from translation posteriorly: *nos* mRNA, slightly enriched in the pole plasm, is translated exclusively at the posterior pole region, and the translated product (NOS protein) diffuses anteriorly, thereby creating an anterior-to-posterior *hb* gradient in the posterior part of the egg. These *Drosophila* embryogenesis mechanisms are now classic textbook examples in embryology, but they represent a highly derived state, presumably reflecting the requirement for rapid development. In most insects, including the flour beetle *Tribolium castaneum* and the cricket *Gryllus bimanculatus*, by contrast, segments are generated sequentially in an anterior-to-posterior order (for review, see Peel *et al*., 2005; Clark *et al*., 2019) with mechanical similarity to those in vertebrate somitogenesis. In *Tribolium*, the involvement of *nos* in segmentation is also reported (also see below) (Schmitt-Engel *et al*., 2012; Rudolf *et al*., 2020). Thus, *hb*/ *nos* interaction may be an evolutionarily conserved function in insects. Posterior to anterior graded distribution of *nos* mRNA is observed in other insects, including hemimetaborans, which possess ancient conserved traits among insects (Lall *et al*., 2003; Dearden, 2006), where this expression of *nos* may function in both axial patterning and PGC formation through its function to maintain naïve cell state.

The silkmoth *Bomyx mori* is a lepidopteran insect, evolutionarily close to diptera, which *Drosophila* belongs to. However, it is quite distinct in terms of segmentation and PGC formation from *Drosophila* and features unobserved in other insects, suggesting a unique evolutionary process (Nakao, 2021). One of such features is the molecular prepattern established during oogenesis, which is required for such embryogenesis processes as segmentation and dorso-ventral pattern formation: the germ anlage is distinguished by the existence of maternally localized structure or molecules (Kobayashi and Miya, 1987; Nakao, 1999, 2008); within the germ anlage, mRNAs of anterior and posterior organizing genes, *orthodenticle* (*otd*) and *caudal* (*cad*), are localized anteriorly and posteriorly, respectively (Nakao, 2012). An inference from such features is that, although it has a large germ anlage, characteristics attributed to long germ insects, including *Drosophila*, the segmentation does not rely on molecular diffusion in such a way as in the case in *Drosophila*, and segmentation proceeds essentially in a cellular environment, not in the syncytium as observed in short germ insects. Indeed, it shares characteristics with both long and short germ insects. Within the large germ anlage, individual segmental anlage can be fate mapped before blastoderm formation and without segment addition zone (growth zone)(Myohara, 1994), reminiscent of the case in long germ insects. However, the segmentation proceeds sequentially in AP order as in short germ insects (Nakao, 2010). *Krüppel* (*Kr*) RNAi perturbation study also indicates a feature shared with both long and short germ insects: the RNAi embryos not only exhibit segmental deletion corresponding to *Kr* gap domain, as in the case of long germ insects but also truncation of the following segments, as in short germ insects (Nakao, 2012). A clear departure from such categorization is also indicated by the results obtained from *hb* RNAi study in *Bombyx*. Unlike other insects studied so far, *Bombyx hb* RNAi does not result in segmental deletion (Liu and Kaufman, 2004; Mito *et al*., 2005; He *et al*., 2006; Marques-Souza *et al*., 2008). Instead, it leads to supernumerary segment formation (Nakao, 2016), although this phenotype might be masked in the RNAi study of short germ insects by the truncation of posterior segments. Remarkable features in *Bombyx* are also observed in PGC formation. Although unlike *Drosophila*, it does not possess structurally recognizable germplasm within newly deposited eggs, *Bombyx* PGCs are detected morphologically and molecularly soon after blastoderm formation, at relatively an early stage as compared to insects that take the inductive mode of PGC formation, where PGCs first appear and are morphologically recognized within the mesoderm of abdominal segments (Miya, 1958; Nakao, 1999; Nakao *et al*., 2006). However, a study of the expression of four *nos* genes (*nosM*, N, O, and P) in *Bombyx* revealed that maternal mRNA for *nosO* appears to be localized within the mesoderm anlage of freshly laid eggs in a pattern that largely corresponds to the site where future PGCs appear (Nakao *et al*., 2008) in the posterior ventral part of the germ anlage;if it truly represents germplasm, it is a unique germplasm localization pattern, indicating a novel inheritance mechanism thus far not seen outside lepidopterans. *nosO* genome editing was then conducted to examine the loss of function phenotype (Nakao and Takasu, 2019). Contrary to the expectations, *nosO* null strains preserved fertility, albeit reduced as oogenesis was severely compromised. Elimination of *nosO* maternal mRNA did not detectably affect fertility, although it appeared to have some responsibilities with the early expression of *BmVLG* (*Bombyx vas*), which marks the earliest stage *Bombyx* PGCs. These results suggested that other *nos* genes act redundantly to specify PGCs in *Bombyx*. Here to pursue this possibility, we examined the roles of other *nos* genes in the PGC formation process.

## Results

### nosO and nosP double RNAi results in abnormalities in both male and female germ line

Of the four *nos* genes, besides *nosO*, mRNAs for *nosM* and N are also maternally provided as described above. While *nosM* mRNA does not show any localization and appears uniformly present within the eggs, *nosN* mRNA exhibits restriction to germ anlage, within which it exists uniformly (Nakao *et al*., 2008). Delving more into the possibility of maternal PGC specification mechanisms, we examined what could result in germ cell formation when the activities of these genes, together with *nosO*, are perturbed during embryogenesis. For this purpose, embryonic RNAi was done, and the effect was examined by mature oocyte number and morphology of newly emerged insects as described previously. The effectiveness of RNAi is shown by the reduction in the specific *in situ* hybridization signal for each gene (Fig.1).

**Fig. 1.**
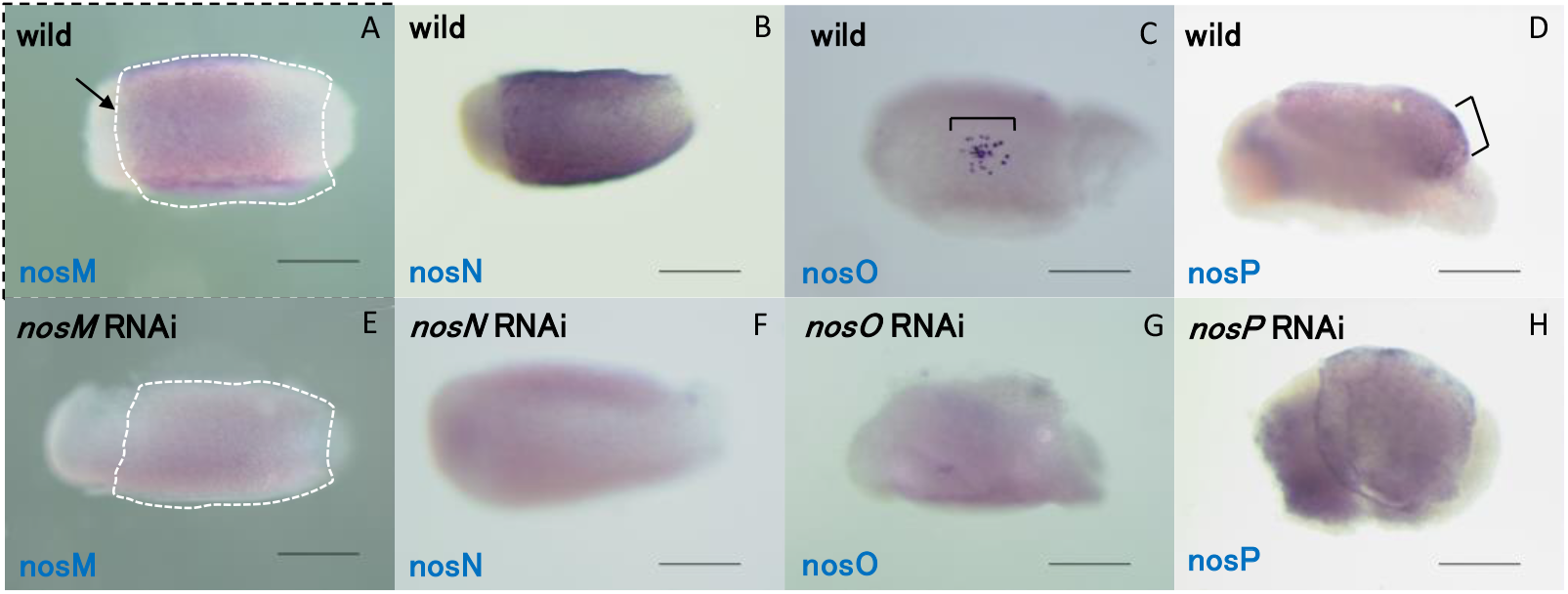
Effectiveness of RNAi targeting *Bombyx nos* genes. RNAi induced degradation of target mRNAs were examined by *in situ* hybridization. Wild-type expression of *nosM* (A), *nosN* (B), *nosO* (C) and *nosP* (D) at 14 h after egg laying (AEL) embryos. Expression of respective genes for *nosM* (E), *nosN* (F), *nosO* (G) and *nosP* (H) RNAi-treated embryos at 14 h AEL. Inside the white dotted lines in (A) and (E) approximate the position of the germ anlage. In (A), *nosM* expressed region is weak but the demarcation of the expressed region and unexpressed part that corresponds to the border of germ anlage and extraembryonic region, is clearly recognizable at the anterior (arrow). For *nosN*, strong expression in (B) is not observed in (F). For *nosO* and *nosP*, typical expression pattern of respective genes (indicated by brackets in (C) and (D); see text, Nakao, 2010) is not observed in treated embryos ((G) and (H), respectively).

When either *nosM* or *nosN* activities are reduced or reduced in combination with *nosO* reduction (*nosM/O, nosN/O*), no effects are observed (data not shown; Fig.2C, D). Moreover, triple reduction of *nosM, N*, and *O* did not detectably affect germ cell formation (data not shown). If *nos* genes are essential in PGC formation as demonstrated in other systems, these results suggest that although we could not rule out the possibility that maternally provided proteins, if in existence, might affect this process, at least *de novo nos* activities from maternal transcript from these genes are not essential in PGC formation in *Bombyx*.

**Fig. 2.**
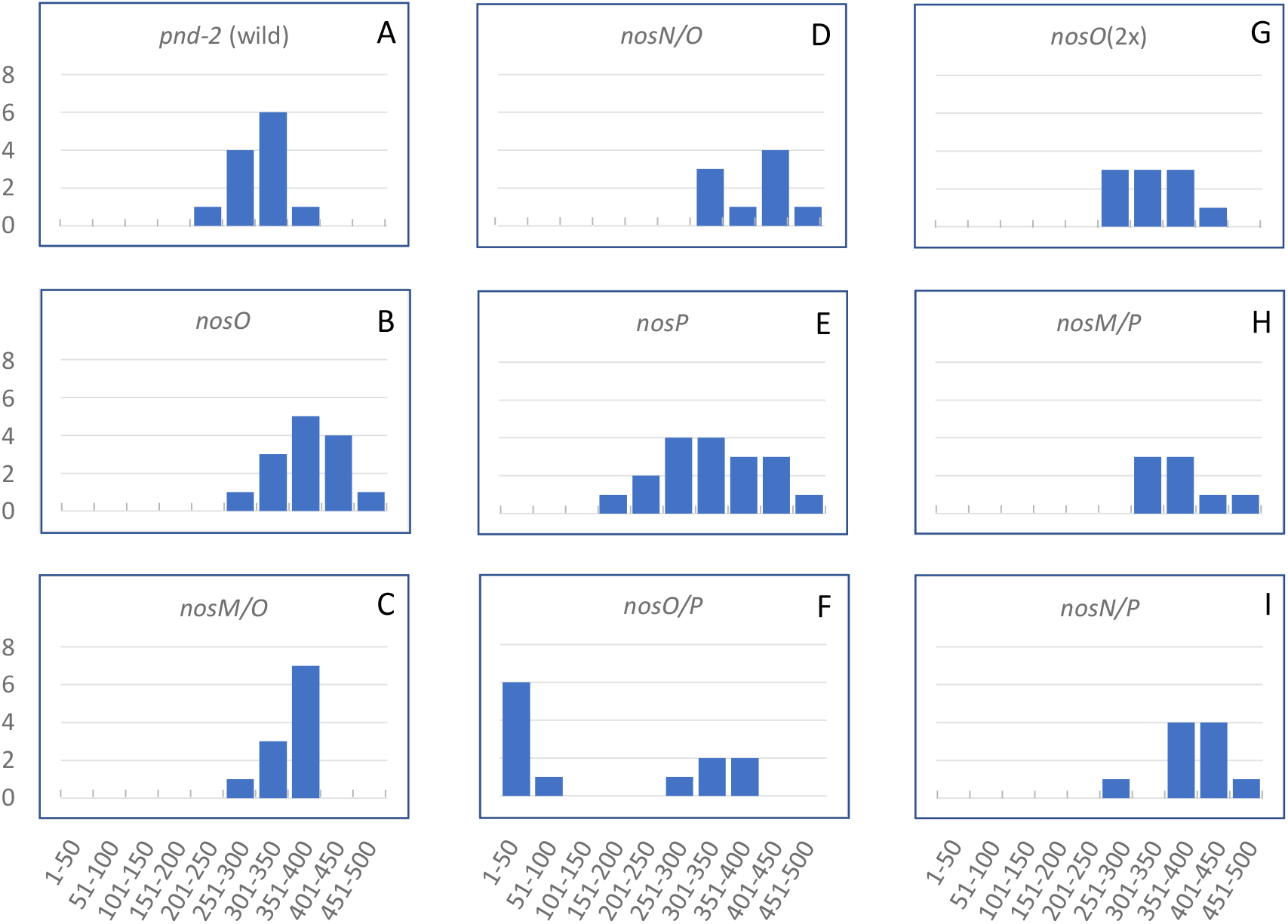
RNAi effect on mature oocyte number in treated female moths. The number of mature oocytes in the control and the RNAi-treated moths were counted, and the number of individuals allocated to each group was plotted.

In contrast to these *nos* genes, *nosP* appears to be expressed exclusively zygotically: its expression begins after germ band formation in its posterior part; as the segmentation proceeds, the expression domain retracts posteriorly following the posterior retraction of PGCs. Indeed, during this retraction process, the *nosP* expression domain appears to continuously cover the region where PGCs exist (Nakao et al., 2008), suggesting that *nosP* may have a role in PGC formation/maintenance. To investigate this possibility, we next examined the phenotype of *nosP* RNAi embryos. The individuals treated for *nosP* RNAi alone exhibited no obvious difference from the untreated ones (Fig. 2E, compare with Fig. 2A). However, when combined with *nosO* RNAi (*nosO/P*), the resulting moths exhibited marked reduction in the oocyte number (Fig. 2F; Fig.3B, compare with Fig.3A), together with aberrant morphology in some remaining oocytes. In most severe cases, the treated female moths had only a few oocytes (Fig. 3B). Also, the testis of *nosO*/*P* treated moths often showed partly transparent appearance (Fig.3D, compared with Fig.3C). To examine the fertility of these male moths, they were crossed with wild-type females. The resulting female moths laid several-fold reduced number of eggs as compared to control (wild vs. wild) crosses, and none of these eggs showed signs of development, while the control eggs were normally developed (Fig. 4). These results indicate that the treated males are sterile. The specificity of this RNAi to both *nosO* and *nosP* genes was confirmed by examining the effect of independently prepared dsRNAs targeting the non-overlapping sequence of each gene. As described above, combinatorial RNAi of *nosO* with either *nosM, nosN*, or *nosM* and *N* did not show any detectable anomalies. The same was true for *nosO* RNAi of double strength (*nosO* (2x)) (Fig. 2G). The insects with a combined reduction of *nosP* with either *nosM* (*nosM/P*), *nosN* (*nosN/P*), or *nosM* and *N* also had no detectable developmental defects (Fig. 2H, I; data not shown).

**Fig. 3.**
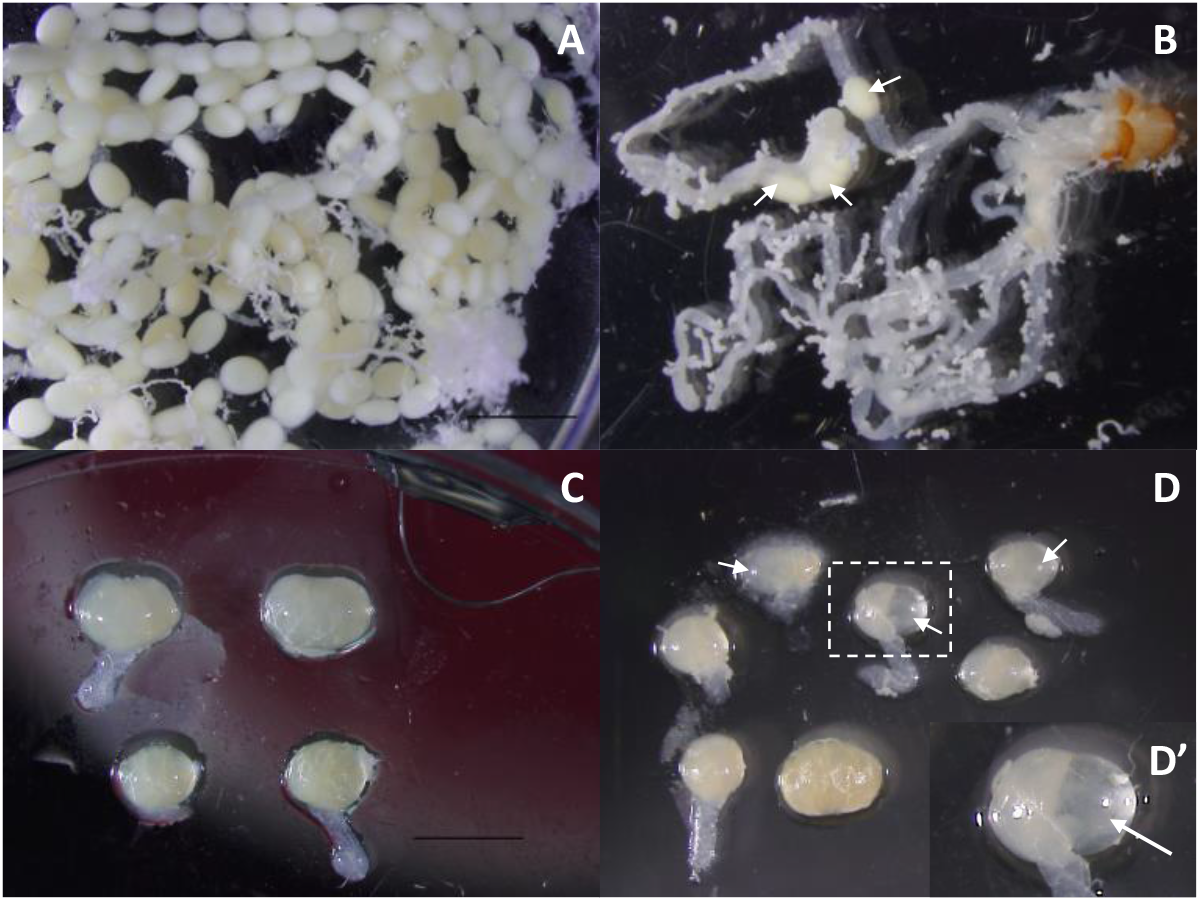
*nosO*/*P* double RNAi effect on gonad formation. (A)(B) Ovaries of wild (A) and RNAi-treated (B) moths. Wild-type ovarioles are full of mature oocytes, whereas the ovarioles of the RNAi-treated moth are basically empty except for one ovariole containing a few developed oocytes (arrows). This RNAi-treated sample was an example of the most severe phenotype. (C)(D) Some examples of testis of wild-type (C) and RNAi-treated (D) moths. In the testis of RNAi-treated moths, the transparent part is often observed (arrows in (D), (D’): higher magnification view of dotted square part).

**Fig. 4.**
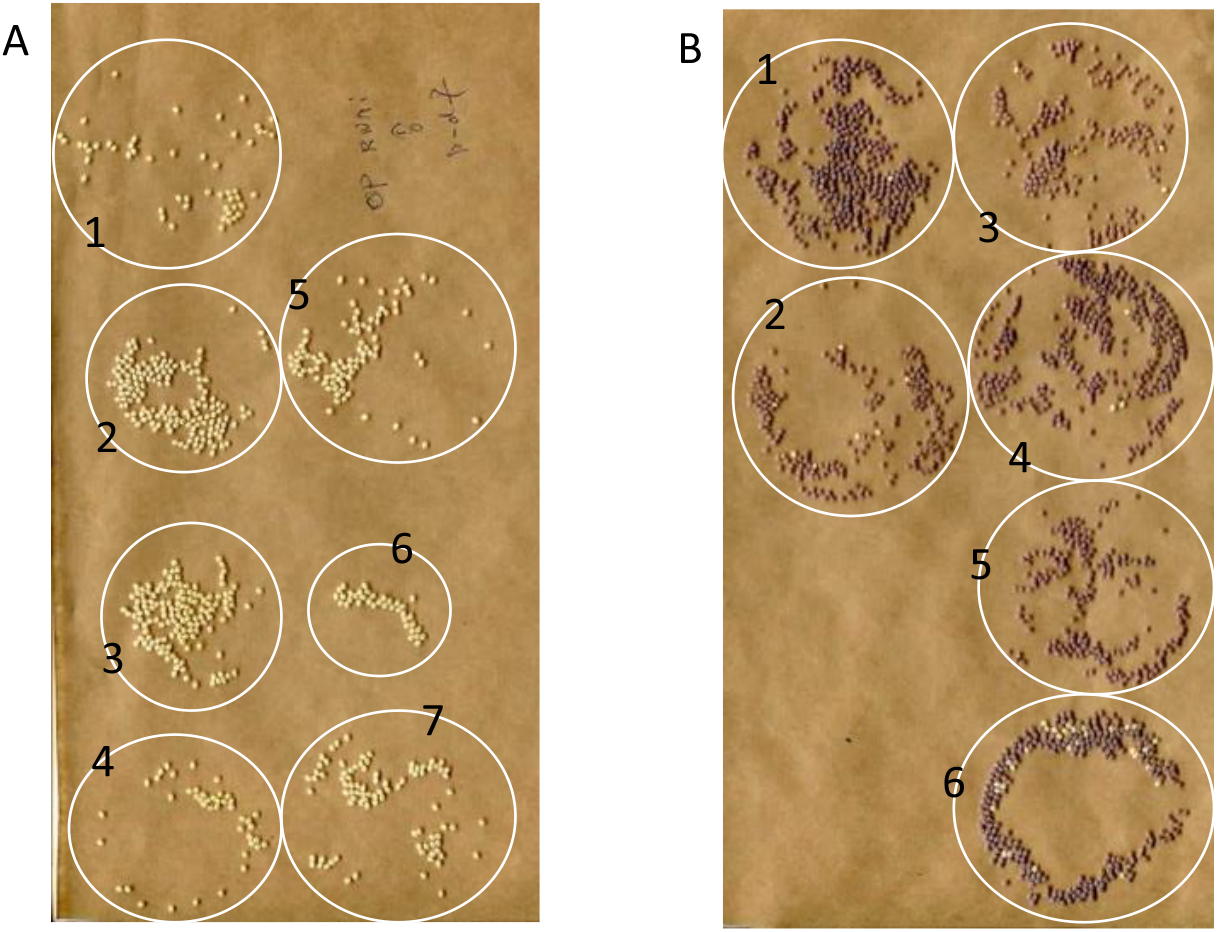
Examination of the fertility of *nosO*/*P* RNAi-treated male moths. Eggs laid by female moths mated with RNAi-treated male moths (eggs laid by each female moth are circled numbered in A), and those with wild-type male moths (circled numbered in B) are shown. Each circle in the photographs indicates eggs deposited by single female moth. Most eggs in B are colored, which indicates the occurrence of normal development, whereas in A, all the eggs lack coloration, indicating the absence of egg activation. For details, see text.

### nosP genome editing produced sterile moths

To confirm and extend the RNAi study in the previous section, transcription-activator-like effector nuclease (TALEN)-mediated *nosP* genome editing was conducted to make knock-out (KO) alleles. Two strategies were employed for their production: introduction of point mutation in the 5′-region of the amino acid coding sequence (CDS) and the removal of a large part of CDS by cutting at two target sites (Fig. 5A). The resulting frameshift mutation in the former strategy was expected to induce nonsense-mediated mRNA decay and produce null mutants.

**Fig. 5.**
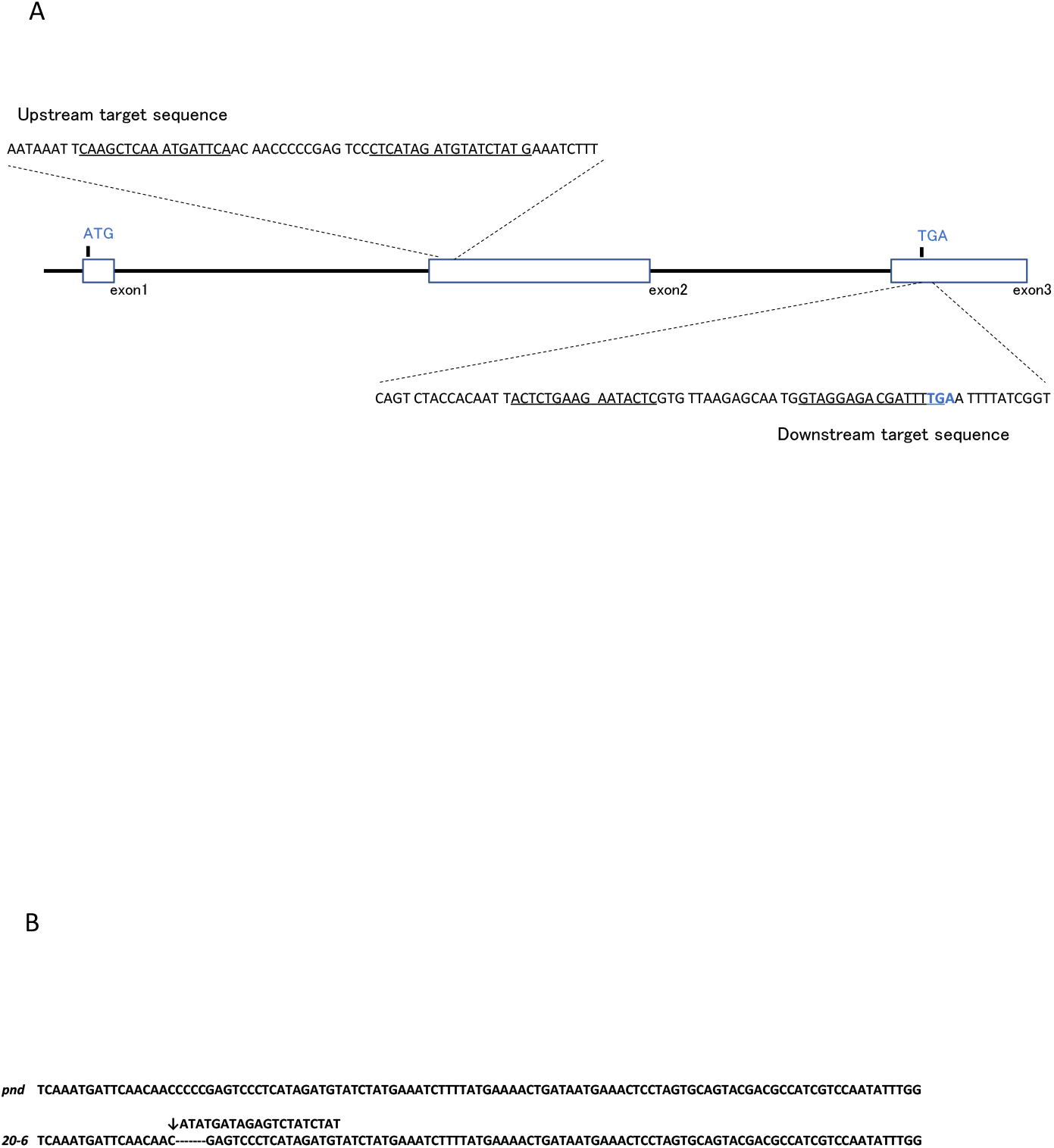
(A) Genomic organization of *nosP* locus. The nucleotide sequences in the vicinity of upstream and downstream TALEN target sites are shown. TALEN recognition sequences are underlined. (B) Nucleotide sequence of an edited allele used for phenotypic analyses. See text for details.

The editing procedure was described previously, and two lines harboring *nosP*-edited allele corresponding to different procedures described above were established. Using these lines, phenotype analyses were conducted. Essentially, the same results were obtained for the resulting edited individuals irrespective of these two strategies.

Unexpectedly, *nosP*(-/-)homozygotic moths were sterile. While testis was not found in males (Fig. 6B, compare with Fig. 6A), female moths were almost devoid of oocytes, i.e., almost all the ovarioles were empty (Fig. 6D, compare with Fig. 6C). However, curiously, a few developed oocytes were almost always observed in the resulting females. To examine the fertility of the edited males, homozygous *nosP*-KO–males were mated with wild-type females, and the resultant eggs were allowed to develop (see **Materials and Methods**). Similar to the case observed in the RNAi experiment, fewer eggs were laid, and they did not hatch, while the control crosses produced a normal number of viable eggs (data not shown). In another experiment, wild-type female moths already subjected to mating with *nosP*-edited males and subsequent egg deposition were thereafter crossed with wild-type male moths and eggs were laid. While a large number of viable eggs were laid after subsequent crossing with wild-type males, very reduced number eggs, all of which were sterile, were laid after prior crossings with *nosP*-edited males (Fig. 7). This not only confirmed the ability of these females to produce viable eggs but also indicated that the functional sperm production ability of males somehow leads to the stimulation of egg laying. The cause of this phenomenon is unknown, but it could be due to a direct effect of the existence of (functional) sperm or some indirect effect of the presence of some stimulating substance(s) or function, production, or function of which is affected by the process of normal sperm generation.

**Fig. 6.**
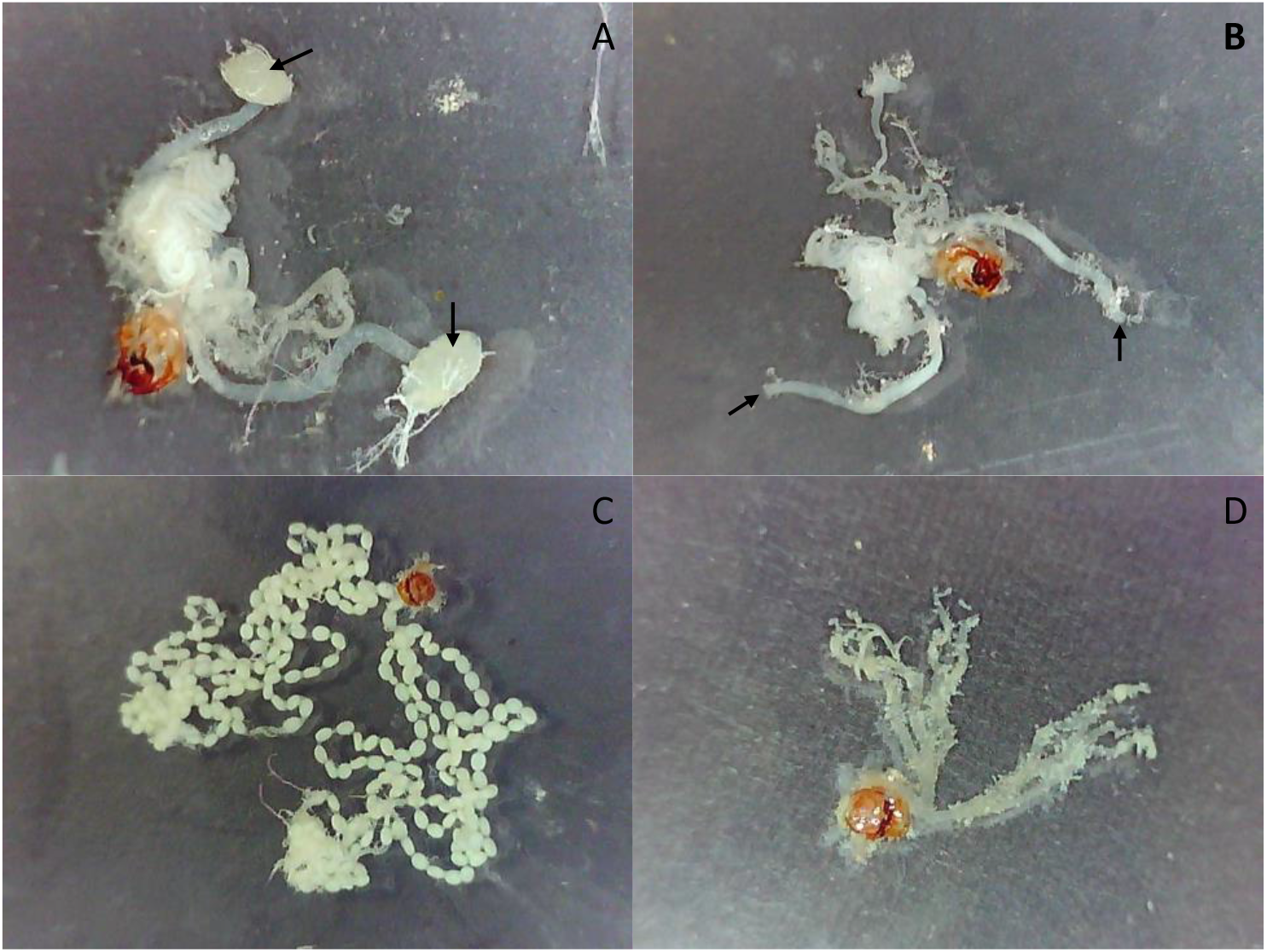
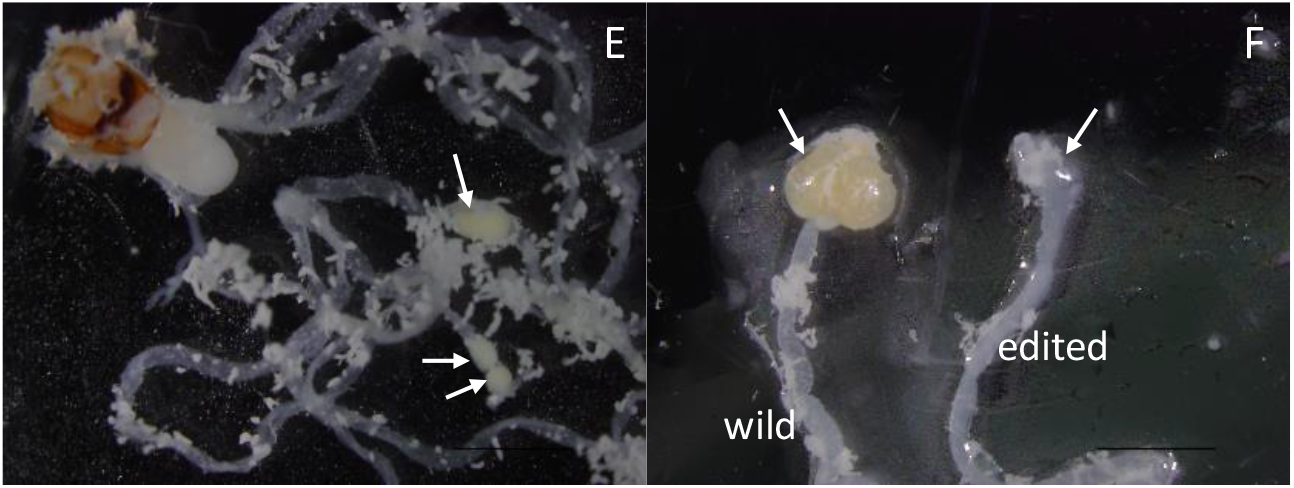
Genital organ morphologies of control male (A), female (C), *nosP*-edited male (B), and female (D) moths. The organ from the edited male lacks testis, which is present in wild-type male (arrows in A, whereas arrows in B indicate where testis should be attached). The organ from the edited female comprises empty ovarioles (D), whereas ovarioles from the wild females are full of mature oocytes(C). Interestingly, edited female individuals almost always contain a few developed oocytes (arrows in E). (F) is the magnified view of the distal end of *vas* deferens where the testis (left arrow) is normally attached in wild-type (left-wild) with the right(edited) showing its absence (right arrow).

**Fig. 7.**
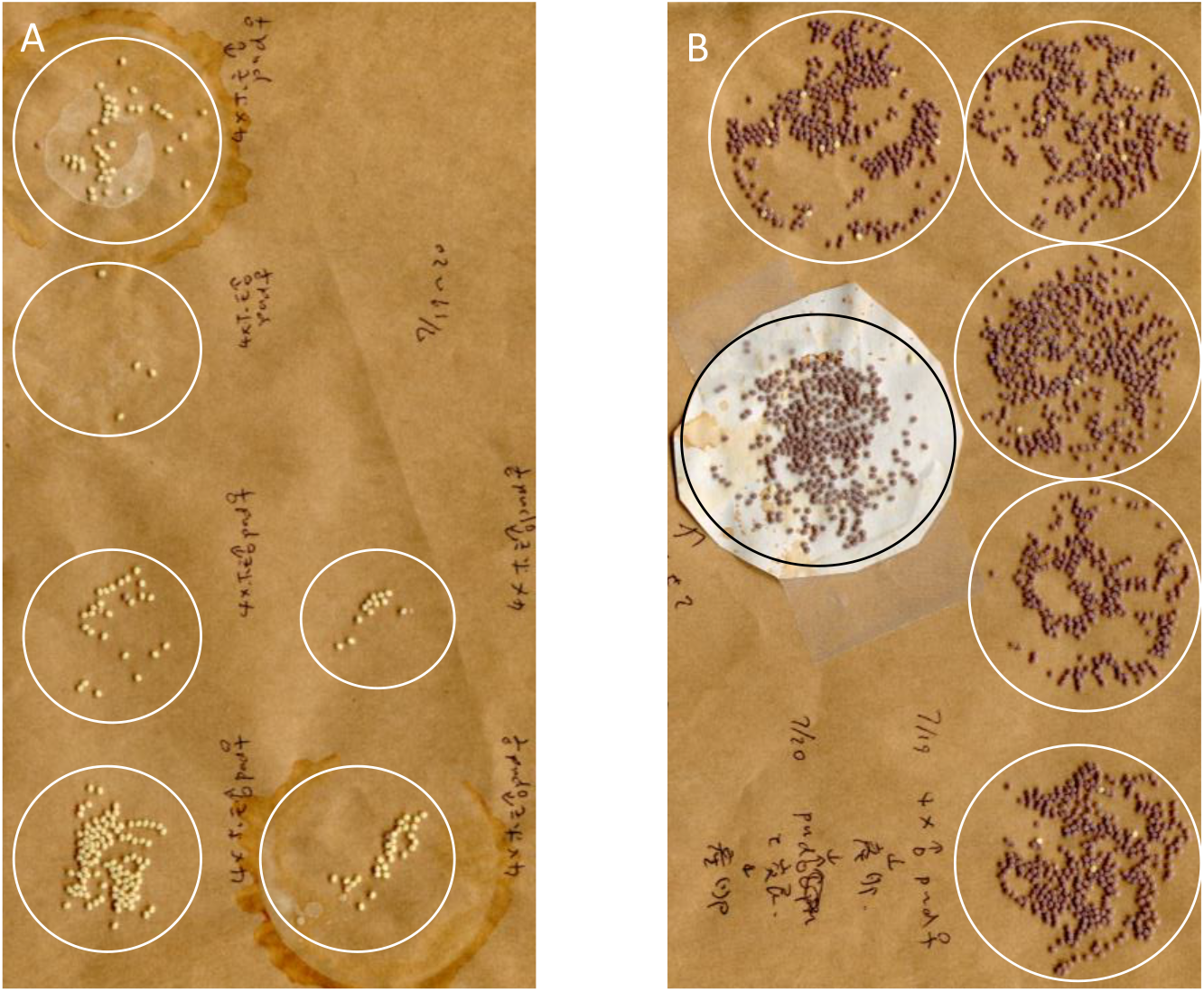
Loss of fertility of *nosP*–KO males and induction of spawning activity by copulation with wild-type male insect. Eggs were first spawned by female moths after mating with *nosP*-edited males (A, total of six pairs). Subsequently, the same females were mated with wild-type males and made them spawn eggs (B). Each circle in the photographs indicates eggs deposited by a single female moth as in Fig.4. As shown, while crosses with edited males produced a small number of sterile (uncolored) eggs, subsequent crosses with edited males produced a large number of fertilized (indicated by purple coloration) eggs, which suggest that copulation with wild-type males somehow stimulates spawning activity, which is absent in *nosP*-edited males.

### Does Bombyx not require nos gene activities in AP patterning?

The study described above indicated that *nosP* gene does not appear to have AP patterning function because the external morphology of *nosP*-KO insects appeared normal (data not shown). This was unexpected since AP patterning functions are considered conserved in insects (see **Introduction**) and *Bombyx nosP* is the only gene expected to be expressed in the posterior restricted manner among the four *Bombyx nos* (see **Discussion**). This raised the possibility that *nos* gene activities are not essential in *Bombyx* embryonic AP patterning. Indeed, no marked difference in embryogenesis was discerned in the moths that have subjected to RNAi procedure for all four *nos* genes as compared to the cases in other RNAi experiments described above: the ratio of eggs that resulted in successful embryogenesis after dsRNA injection did not appear to show marked difference. This is consistent with the fact that healthy female moths were obtained after RNAi against four *nos* genes except for showing a severe reduction in mature oocyte number, which testified that the RNAi procedure is functional despite their high dsRNA load. However, these moths could have somehow alleviated the morphological effect of the other *nos* genes, if any, that resulted in the completion of embryogenesis, leading to the successful hatching and the growth to adulthood, whereas other affected embryos might have died with such an effect unrecognized. To obtain more unbiased result, embryogenesis was examined for eggs with features of successful dsRNA injection (see **Materials and Methods**) after dsRNA injection more directly by dissection 5 days after egg laying, when the basic larval features became apparent. About one-third of both control (enhanced green fluorescent protein dsRNA [12 μg/μl] injected-) and *nos* dsRNA (3 μg/μl for each: total of 12 μg/μl) injected embryos showed normal appearance (n=6/15 for control and n=5/16 for *nos* injected). The other embryos from the control showed deficiencies in the anterior (head) part of the embryos. The cause of this phenotype is unclear, but it might be due to some mechanical stress caused by the deformation of the eggshell as an effect of injection or might be caused by the injection of large amount of dsRNA, or some other reasons. From the *nos* RNAi-treated embryos, embryos showing some anomalies in the posterior regions of the embryos were also observed (n=6/15). The significance of this finding is unclear. For these embryos, some perturbation in the morphogenetic process appears to have occurred after normal patterning, whether this is a *nos* RNAi specific effect also remains to be clarified. This high rate occurrence of normal embryos appears to be significant considering that, in our hands, usually, a specific phenotype had occurred at very high penetrance (≥90%) after RNAi of developmentally essential genes (*otd, cad, Kr*, pair-rule genes; Nakao, 2010, 2012). These results suggest that the function of *nos* genes in *Bombyx*, if any, does not consist of the main part of the embryonic patterning, but is conditionally evoked for its robustness.

## Discussion

### Bombyx nos function in germ cell formation/maintenance

In the previous study, the functional analysis of *nosO* by embryonic RNAi and gene editing was performed since its expression pattern suggests its critical roles in PGC specification/maintenance. We found severe abnormalities during oogenesis, and some hint at its involvement in PGC specification/maintenance in *nosO* KO’s. However, albeit severely reduced, the fecundity was not completely eliminated, which made it possible to establish *nosO* KO lines. Indeed, the plausible effect on PGC specification/maintenance, i.e., generating female insects with a severely reduced number of oocytes, was only rarely observed in *nosO*–KO insects. Considering the *nos* gene plays essential roles in the maintenance of PGCs in *Drosophila* or in mice, these observations suggested the possibility that the other *nos* genes complement the function of *nosO*. This study addressed this possibility and identified such a function in *nosP* during embryogenesis, the functional redundancy between *nosO* and *nosP*: while individual RNAi study against these genes did not lead to the reduction in the oocyte cell number, the simultaneous RNAi did. In the most severe cases, the female moths were almost devoid of oocyte, and similarly treated male moths lost fecundity. This RNAi, together with *nosP* KO results suggest that the RNAi exerts its effect on the embryonic expression of *nosO* and *nosP*, and that these genes are involved in PGC formation/maintenance. Although the previous study has seen considerable cases of oocyte number reduction in the *nosO* RNAi experiment, such an effect was not observed in this study. The reason for this discrepancy is unclear. Possible explanations for this could be either that the characteristics of silkmoth strain might have changed during successive rearing or that experimental conditions might not have been identical. For the latter, the embryonic RNAi experiments as conducted in this study have at present some uncontrollable elements in our hands, and the hatching rate after injection differs greatly between experiments. Indeed, this series of RNAi experiments was generally conducted under higher hatching rate after injection as compared to the previous study.

As described in the **Introduction**, metazoan PGCs are known to be specified by zygotic induction or inheritance, which is derived from zygotic induction. Because PGC specification and embryonic axial patterning mechanisms appear intimately linked, it is highly likely that the inheritance mechanism co-evolved with the evolution of the axial patterning mechanisms. Presently, the molecular definition of PGCs is ambiguous, but the main functions are considered to be the preservation of the genome integrity and totipotency. For this purpose, mechanisms appear to have been evolved during PGC specification process to counteract the insults inflicted on PGC precursors accompanied by the progression of axial patterning. These mechanisms may differ depending on the axial patterning mechanisms, which vary greatly between insects, i.e., there may be various molecular paths toward PGC specification. This can be seen in the requirement of *nos* gene function in PGC specification/maintenance. In *Drosophila*, slightly posterior-pole enriched maternal *nos* mRNA is exclusively translated at the posterior pole, and translated protein either diffuses within the syncytium anteriorly to repress *hb* mRNA or are sequestered directly within pole cells at its formation; in the pole cells (PGCs) thereafter, NOS protein represses somatic gene expression to maintain an undifferentiated state (Kobayashi *et al*., 1996). In the honeybee *Apis melifera*, however, *nos* gene function may not be required in the early embryonic PGC specification/maintenance processes (Dearden, 2006). In *Bombyx*, this RNAi study indicated the absolute requirement of nos function in PGC specification/maintenance and clarified the redundancy of *nosO* and *nosP* for this function. Despite such a redundancy, their spatial expression patterns differ; while *nosO* mRNA enriches in PGCs, *nosP* appears to be expressed uniformly in the posterior part of the embryos irrespective of cell type. The previous study could not clarify the function of maternal *nosO*, which is likely mediated through the localized mRNA that regulates the expression *BmVLG* along with its zygotic activities (Nakao and Takasu, 2019). The functional redundancy described above suggests that the maternal *nosO* mRNA contributes to robust PGC formation and is indeed involved in the inheritance mechanisms: without *nosO*, the PGC formation would be vulnerable to perturbations. This feature of *nosO* suggests a situation similar to *Drosophila*, although functional requirement during early embryonic stages appears limited, as suggested from the previous study (Nakao and Takasu, 2019). However, *nosP* influence on PGC formation is reminiscent of zygotic induction in that it is expressed broadly in the posterior of the embryos in a non-PGC specific manner and the mRNA expression pattern does not suggest that the gene activity provides a cue for the cells to become PGCs. Recent studies in *Gryllus* clarified some features of zygotic induction mechanisms in insects, in which similarities to vertebrate PGC specification mechanisms have been uncovered, such as the involvement of *Blimp1* and TGF-β(BMP)signaling (Donoughe, *et al*., 2014; Nakamura and Extavour, 2016). These and other studies indicate that PGC specification occurs progressively (Ewen-Campen *et al*., 2013; Barnett *et al*., 2019): in *Gryllus, HOX* genes are revealed to limit PGCs to specific abdominal segments. Likewise, the *nosP* may function in limiting PGCs to its expressed zone. These suggest such *nosP* function, which acts either cell-autonomously or cell-non-autonomously, or both, might be a remnant of the ancestral zygotic mechanisms, where *nos* function might be required for the specification of PGCs. This idea is consistent with both the evidence that *nosP* preserved ancestral characteristics among the four *nos* genes and phylogenetically close to mammalian *nos* genes (De Keuckelaere *et al*., 2018), and with the result in this study that *nosP* KO leads to virtually complete sterity, which suggests that *nosP* plays central roles in germ cell functioning. *nosO*, however, may have later evolved to gain specialized function in PGC specification/maintenance with the evolution of axial patterning mechanisms by acting as a player in the inheritance mechanisms and contributed to the early PGC specification seen in *Bombyx*. The previous study suggests the existence of maternal cue colocalized with *nosO* mRNA: even when maternally localized *nosO* mRNA is eliminated, PGCs as examined by *BmVLG* (*Bombyx vas* homolog) still appear in the similar location and at a similar time as in the wild-type (Nakao and Takasu, 2018) without detectable localized zygotic *nosO* mRNA, suggesting that this feature represents an evolutionary path toward its unique PGC specification mechanisms.

In *Gryllus*, the mesodermal origin of PGCs is also suggested; RNAi knockdown of *twist*, a gene essential for mesoderm formation, led to the elimination of PGCs. Whereas in *Bombyx* eggs, maternal *nosO* mRNA appears to be localized in the mesodermal anlage (Nakao *et al*., 2008; Nakao, 2010), which suggests that *Bombyx* PGCs arise from cell population destined to be part of mesoderm in the absence of such a maternal cue. From this, we can speculate that these two species have in common some mechanisms in PGC formation, which may indicate an evolutionary relationship or deep homology. This apparently shared feature may be related to the fact that the germ anlage of these insects occur as a ventral anlage without covering the posterior pole (Pechmann *et al*., 2021; Nakao, 2021). However, presently, the detailed ontogeny of *Gryllus* PGCs is unclear and there remains the possibility that the RNAi result described in the *Gryllus* study implies the necessity of mesoderm for the maintenance of PGCs, and not PGCs mesodermal origin. In contrast to the case in *Bomyx*, however, available information does not indicate the provision of a maternal cue in *Gryllus*: *vas* and *piwi* parental and embryonic RNAi do not affect PGC formation, consistent with the view that zygotic induction mechanisms operate in this species. These suggest that the ancient mechanisms of PGC formation as seen in *Gryllus* are modulated by the existence of a maternal/preexistent zygotic cue in *Bombyx*; such a cue (or bias) to be PGCs might have substituted for the stochastic process possibly occurring in ancestral zygotic induction to, for instance, make *Bombyx* PGC formation robust and this function of maternal cue could be responsible for the earlier appearance of PGCs in *Bombyx* compared to such insects as *Gryllus* that takes induction mode of PGC formation. However, because how PGCs are formed in *nosO* KO’s is still unknown, such as their ontogeny, for instance, the possibility that in *Bombyx*, both ancient (zygotic induction) and derived (inheritance) mode of PGC formation operate in parallel, i.e., the dual ontogeny, cannot be ruled out. Considering the importance of *nos* gene in this context, it is also important to know the role of *nos* gene in *Gryllus* PGC formation, which is currently unavailable. From this perspective, it is interesting to know whether *nosO* and *nosP* have different targets and, if this is the case, their identity. Such information could provide an example of molecular mechanisms toward inheritance. In *Drosophila*, a classic example of harboring inheritance mechanisms in PGC specification, TGF signaling after fertilization was recently shown to be involved in the pole cell formation process by modulating the action of pole plasm, indicating the involvement of zygotic mechanisms in PGC specification. Thus, the cases of involvement of both mechanisms of PGC formation in one organism may be widespread as seen in an example in this study, and such mechanisms may be important for the integrity of organismal development. A recent review on insect PGC formation mechanisms suggests the liability of PGC specification mechanisms. Based on the lack of *oskar* gene in *Bombyx*, its PGC formation mechanisms have been suggested as an example of returning to ancestral zygotic mechanisms (Lynch *et al*., 2011). The results obtained in this study do not clarify whether this is indeed the case. However, it might be possible that the simultaneous existence of both mechanisms contributes to such liabilities.

### nos function in early Bombyx embryogenesis

As described above, inheritance mechanisms appear to have evolved to counteract the effect of the somatic developmental program accompanied by the evolutionary change in axial patterning mechanism. For example, *Drosophila* and *Nasonia* developed mechanisms of segregating PGCs as pole cells located at the posterior pole before the beginning of the somatic program. This strategy is effective because, in these insects, axial patterning occurs in an environment where molecular diffusion is allowed at the syncytium blastoderm stage after pole cell segregation. By contrast, *Bombyx* PGCs appear relatively late after blastoderm formation among cells undergoing somatic development. *Bombyx* strategy for early development is unique in that the periplasm of newly deposited eggs has maternally established localized distribution of mRNA for organizing molecules specifying early embryonic development, a condition in *Drosophila* that appears to be largely corresponds to that attained after diffusion of these molecules in the syncytium. This suggests the possibility that cells that eventually become PGCs receive positional information similarly dictating early embryonic development to cells destined to soma. Maternally localized *nosO* mRNA could contribute to counteracting this effect toward somatic development, although this function could not be experimentally detected in the previous studies.

With their early embryonic expression patterns, *nosM* and *nosN* functions may also reflect the unique early embryogenesis feature of *Bombyx*. As a translational regulator, *nosM*, which appears to be distributed within the eggs uniformly, might be involved in the initiation of embryonic development or an event as zygotic gene activation, or *nosN*, which is expressed uniformly within the germ anlage, might have a function in fine tuning the timing of translation of localized transcript within the germ anlage, which might ensure the function of the pre-established boundary between embryonic vs. extraembryonic region.

### Is nos involved in embryonic AP pattern formation in Bombyx?

Although validated cases are scarce, another possibly conserved function of *nos* other than those in germ cells in insects is their involvement in embryonic AP patterning through repression of *hb* translation. The function for abdomen development in *Drosophila* is well-known, as described in the **Introduction**. In *Tribolium*, RNAi perturbation of *nos* function is reported to result in developmental arrest by affecting the process of posterior segmentation and in another study, in the acceleration of blastodermal AP patterning process (Schmitt-Engel *et al*., 2012; Rudolf *et al*., 2020). Such functions of *nos* may reflect the posterior expression of *nos* genes observed in some insects (Lall *et al*., 2003). Of the four *nos* genes identified in *Bombyx*, only *nosP* mRNA exhibited a posterior expression pattern. Therefore, it was rather surprising that *nosP* KOs develop normally because it was expected that *nosP* function perturbation leads to AP patterning defects. Additionally, the fact that embryonic RNAi targeted at all four *nos* genes simultaneously did not result in marked difference in embryogenesis from wild-type might indicate that *nos* gene functions are dispensable for AP patterning in laboratory environment. These are, however, reasonable if the prime target of *nos* in AP patterning is *hb* and considering the peculiarity of *Bombyx* AP patterning mechanisms. Of the examined insects for *hb* function by RNAi studies, *Bombyx* is exceptional in that it did not lead to posterior truncation phenotype; instead, it leads to supernumerary segment formation as described in **Introduction**, suggesting that, in *Bombyx, hb*/*nos* system for insect AP patterning may not operate: *Bombyx* would have developed a means to restrict hb expression that does not rely on *nos* functions. Since *hb* intersect with both AP patterning and PGC formation, changes observed in *hb* expression/function could have significant implications in considering the evolution of unique features of *Bombyx* embryogenesis.

## Materials and Methods

### Silkworm strains, rearing, and development

*Bombyx mori* strain pnd-2 used in this study were reared on an artificial diet (Nippon Nosanko) at 28°C. For a general description of early *Bombyx* development, refer to Nagy *et al*., (1994) and Nakao (2021).

### Embryo fixation, in situ hybridization, and RNAi

Embryo fixation, *in situ* hybridization, and RNAi were performed essentially as previously described (Nakao, 1999, 2012; Nakao *et al*., 2006). In simultaneous RNAi against multiple targets, dsRNA concentration for each target in injection solutions was 3 μg/μl. dsRNAs were prepared using MEGAscripts RNAi Kit (Ambion) exactly as described in the manual. The templates used for *in vitro* transcription were PCR fragments of the corresponding genes, flanked by T7 promoter sequences. The primers used for amplification of those were as follows; *nosM*: 5’-taatacgactcactatagggagagtacgtttcgtttgtcatca-3’, 5’-taatacgactcactatagggagaacactgactccccatttttc-3’; *nosN*: 5’-taatacgactcactatagggagaggagagcaaagagcaacatcttcgt-3, 5’-taatacgactcactatagggagacgacacgtagttgttagcag-3’; *nosO*: 5’-taatacgact-cactatagggagaagtaactaaacgcgcctcga-3’, 5’-taatacgactcactatagggagatcagggtctcattgcgcaca-3’; *nosP*: 5’-taatacgactcactatagggagacaagcattcgatccatcgtg-3’, 5’-taatacgactcactatagggagactgatctgctctctttcgga-3’. Primers for amplification of second non-overlapping *nosO* and *nosP* target; *nosO* (2): 5’-taatacgactcactatagggagaaagtgcagcccaccgaggag-3’, 5’-taatacgactcactatagggagactgttccagggcagcccaaa-3’; *nosP* (2): 5’-taatacgactcactatagggagacttttctatgacatcttcggacttg-3’, 5’-taatacgactcactatagggagattcgttggctttctttgcgg-3’. After injection, irrespective of the injected materials (dsRNA or DNA construct etc.), the eggs with an air bubble or severely recessed by desiccation appear at various (often high) frequencies, and they do not complete embryogenesis. These eggs are not considered as “successful injection” and are omitted from morphological analyses.

### Generation of nosP knockouts

*nosP* KO’s were generated by employing TALEN-mediated genome editing with one or two target sites (Takasu *et al*., 2013). For the former procedure, we set a target site within the second coding exon and expected a pair of TALENs to introduce a frameshift within the coding sequence, resulting in the generation of premature termination codon and degradation of mRNA by nonsense-mediated decay pathway. For the latter, target sites were selected such that a large portion of CDS is removed to eliminate the gene function. Procedures for KO silkworm generation were described previously. In brief, TALEN vectors were constructed by Golden gate assembly (Cermak *et al*., 2011), *in vitro* transcribed using HiScribe T7 ARCA mRNA synthesis kit (New England Biolab) and microinjected into silkworm embryos within X hours after oviposition. The resultant moths (G0) were crossed to the wild-type moths to obtain G1 offspring, from which genomic DNA was extracted in adulthood. Screening for successfully edited silkmoths was conducted by PCR using a pair of primers outside the target site(s) and for the former procedure, and additionally by subsequent sequence analysis. The TALEN target sequences and genomic organization of *nosP* are shown in Fig. 6A. The genome sequence of the edited allele, designated as 20–6, obtained by the former procedure, which was used for subsequent phenotypic analyses, is shown in Fig. 6B compared with wild-type sequence. In this allele, the 4 bp at the center of the TALEN target is substituted by 20 bp insertion sequence.

### Examination of male fertilization ability

Male fertilization ability was measured by examining the phenotype of eggs deposited by females after copulation with RNAi-treated or -edited males and comparing them with control mates using wild-type males. Newly emerged females were used for this study. After a few hours of mating, the male and female moths were separated, and the females were left to lay eggs overnight. Successfully fertilized eggs were either known by the coloration of serosa, which is observable after a few days of egg deposition, or a sign of cuticle development, which is visible through the chorion at later stages.

## Data availability

Not Applicable

